# Arborist: Prioritizing Bulk DNA Inferred Tumor Phylogenies via Low-pass Single-cell DNA Sequencing Data

**DOI:** 10.64898/2026.02.26.707773

**Authors:** Leah L. Weber, Chi Yin Ching, Christopher Ly, Yidan Pan, Yixiao Cheng, Chunxu Gao, Peter Van Loo

## Abstract

Cancer arises from an evolutionary process that can be reconstructed from DNA sequencing and modeled by tumor phylogenies. High coverage bulk DNA sequencing (bulk DNA-seq) is widely available, but tumor phylogeny inference requires deconvolution, often resulting in non-uniqueness in the solution space. Single-cell DNA sequencing (scDNA-seq) holds potential to yield higher resolution tumor phylogenies, but the sparsity of emerging low-pass sequencing technologies poses challenges for the study of single-nucleotide variants. Increasing availability of data sequenced with both modalities provides an opportunity to capitalize on the advantages of these technologies. While inference methods exist for bulk DNA-seq and for low-pass scDNA-seq, no joint inference methods currently exist. As a first step, we propose a method named ARBORIST that prioritizes tumor phylogenies inferred via bulk DNA-seq using low-pass scDNA-seq data. ARBORIST takes as input a candidate set of trees with corresponding SNV clustering, along with variant and total read count data from scDNA-seq and uses variational inference to approximate a lower bound on the marginal likelihood of each tree in the candidate set. On simulated data, matching characteristics of current scDNA-seq data, ARBORIST outperforms both bulk and low-pass single-cell reconstruction methods. On a biological dataset, ARBORIST conclusively resolves the evolutionary relationship between different SNV clusters on a malignant peripheral nerve sheath tumor, which is supported by orthogonal validation via a proxy for copy number. ARBORIST provides a principled framework for integrating bulk DNA-seq and low-pass scDNA-seq data, improving confidence in tumor phylogeny reconstruction.

**Availability:** https://github.com/VanLoo-lab/Arborist

## 1 Introduction

Cancer arises as the result of an evolutionary process, driven by the accumulation of somatic mutations that produce genetically heterogeneous tumors [1]. This evolutionary process is modeled by a tumor phylogeny, which depicts the ordering and branching patterns of these subclone-defining somatic mutations (Fig. 1a). Accurate phylogeny reconstruction from DNA sequencing data deepens our understanding of cancer drivers, clonal trajectories, and therapeutic resistance [2–11].

**Figure 1.**
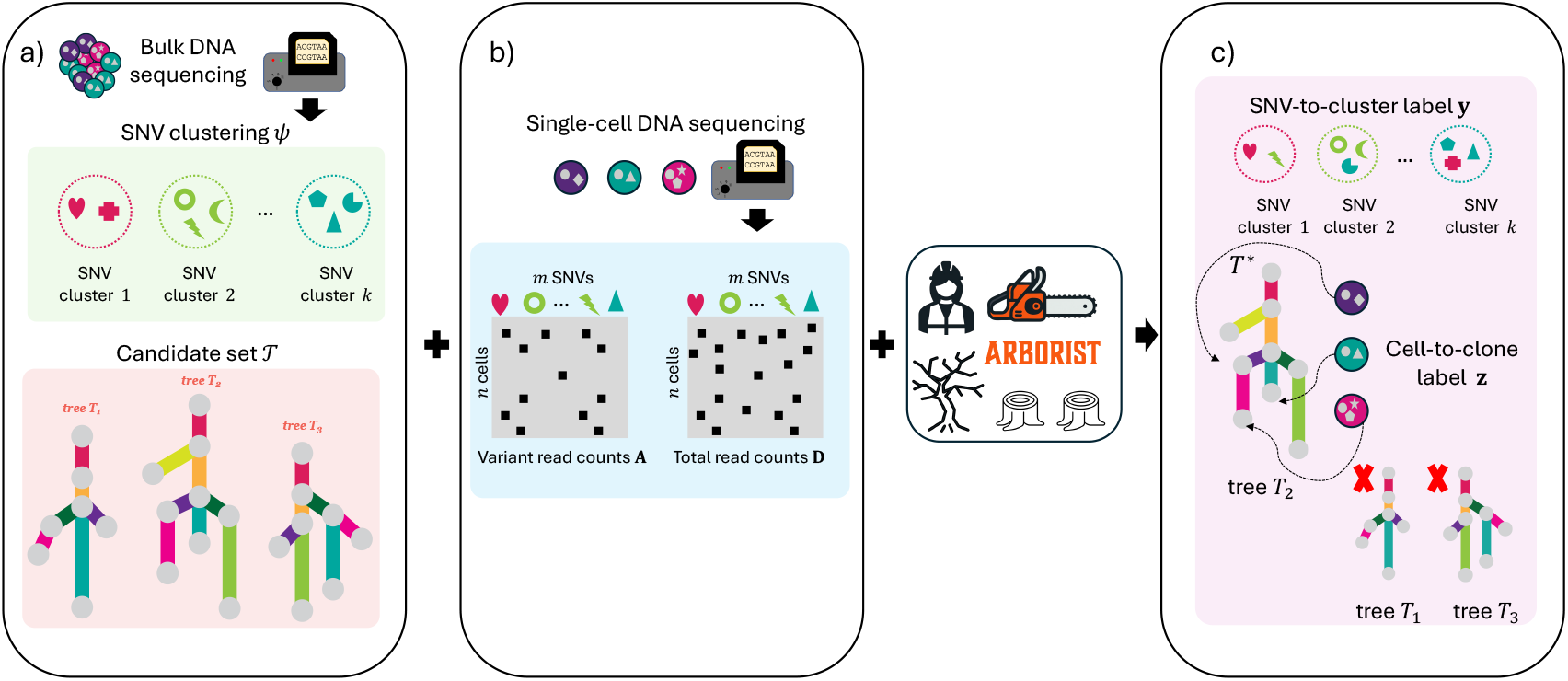
Overview of Arborist. (a) Using existing methods, bulk DNA sequencing data of a heterogeneous tumor is used to infer an initial SNV clustering *ψ* and a candidate set of clonal trees 𝒯. (b) From the *m* SNVs called from the bulk sequencing data, variant **A** and total **D** read counts are generated for each of the *n* single cells. (c) Arborist uses both bulk sequencing and scDNA-seq data to identify the most likely clonal tree *T*^∗^ ∈ 𝒯, as well as the maximum a posteriori (MAP) SNV-to-cluster labels **y** ∈ [*k*]^*m*^ and cell-to-clone labels **z** ∈ [*k* + 1]^*n*^.

To date, whole-genome DNA sequencing data remains a valuable source for tumor phylogeny reconstruction. Besides bulk DNA sequencing, single-cell DNA sequencing (scDNA-seq) is also frequently used. Bulk DNA sequencing has the advantage of being cost-effective for sequencing millions of cells at high coverage, which facilitates the sensitive detection of point mutations, including single-nucleotide variants (SNVs) [12] as well as multi-sample, multi-region, and/or spatial analysis of tumor growth and metastatic patterns [6, 7, 13, 14]. On the other hand, bulk sequencing yields a mixture of cells from a heterogeneous tumor, which requires clonal mixtures to be deconvolved as part of the tumor phylogeny inference problem [15–17]. Deconvolution provides added complexity, resulting in non-uniqueness in the solution space of inferred phylogenies [18]. Despite this challenge, this problem has been well studied and a number of methods exist for subclonal reconstruction and tumor phylogeny inference [15, 16, 19–25]. Consequently, these methods either output a solution space of trees [24] or select a single representative tree [20, 21] even while returning multiple trees. However, many downstream analysis methods, such as the study of metastatic migration or copy number evolution, typically require a single input clone tree [4, 19]. One solution to this challenge is to condense a solution space of clone trees to a single point estimate by consensus tree building methods [26]. However, while a single consensus tree is convenient, it may miss important topological features [27].

In contrast, single-cell DNA sequencing (scDNA-seq) permits high-throughput whole-genome sequencing of individual cells [8, 9, 28]. These technological advancements have led to new insights into intra-tumor heterogeneity and clonal evolution by taking advantage of the uniform sequencing coverage to study copy number alterations (CNAs), which are gains and losses of genomic regions. However, these technologies like DLP+ and ACT, are referred to as low-pass scDNA-seq technologies because the sequencing coverage is extremely sparse, typically around 0.01 × - 0.05 × (Fig. 1b). This has posed significant challenges when studying the evolution of SNVs since many loci of interest within a single-cell may not even be covered by a single read. However, subclonal reconstruction and tumor phylogeny inference methods exist for low-pass scDNA-seq data [29–31], which typically rely on copy number information to overcome the sparsity of coverage.

Given that both types of DNA sequencing have strengths and weaknesses for tumor phylogeny inference, an appropriate strategy to improve accuracy is to utilize both bulk sequencing and scDNA-seq data when available. To date there are two methods, B-SCITE [32] and PhISCS [33] that seek to integrate both data types for DNA sequencing. An additional method called Canopy2 [34] integrates bulk sequencing and single-cell RNA-sequencing data. While these three methods successfully demonstrated that combining bulk and single-cell data is beneficial, one major drawback is that they were designed for early single-cell DNA sequencing technologies that yielded much smaller input sizes in terms of the number of cells and SNVs, typically on the order of hundreds. Therefore, the input sizes of current scDNA-seq technologies, which may yield thousands of cells and tens of thousands of SNVs, pose a challenge for these hybrid tumor phylogeny inference methods.

In this work, we propose a novel approach to integrate bulk sequencing and high-throughput low-pass scDNA-seq data. Our key idea is to build on the many existing methods for bulk tumor phylogeny inference using a two-step process called Arborist. First, we use existing methods to generate a noisy initial SNV clustering and corresponding candidate set of clone trees. Then, given a candidate set of trees and SNV clustering, Arborist utilizes scDNA-seq data to rank each tree in the candidate set by how well it explains the scDNA-seq data (Fig. 1c). In addition to providing a ranking of the candidate set, Arborist makes use of variational inference to yield a mapping of cells to clones and an updated SNV clustering, which is useful for further downstream analyses.

## 2 Materials and methods

### 2.1 Background and problem statement

Suppose we have sequenced *b* bulk samples with bulk sequencing, from which *m* SNVs have been identified. Using existing subclonal reconstruction methods [15, 25, 35], we cluster the *m* SNVs into *k* clusters, i.e., *ψ* : [*m*] → [*k*]. Our goal is to infer a clone tree *T* (defined below) that explains the evolutionary history of the tumor under the infinite sites assumption. Given an SNV clustering *ψ*, we consider a candidate set of 𝒯 clone trees generated by one or more subclonal reconstruction algorithms [16, 20, 21, 24].

#### Definition 1

A rooted labeled tree *T* is a *clone tree* if there exists a bijection *η* : [*k*] → *V* (*T*) between the set of SNV clusters and the nodes of tree *T*.

The bijection *η* defines the labeling of *T*. For each cluster *q* ∈ [*k*], we refer to node *η*(*q*) as clone *q*. Clone *q* is defined by the set of SNVs assigned to all clusters on the path from the root of *T* to node *η*(*q*).

Suppose in addition to bulk sequencing data, we have also sequenced *n* cells with low-pass single-cell sequencing (scDNA-seq) [8, 9, 28] and derived variant 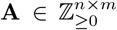 and total 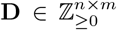 read counts for each of *n* cells and *m* SNVs. Our goal is to identify the clone tree *T* among the candidate set 𝒯 with the highest posterior probability *P* (*T* | **A, D**, *ψ*) given the observed scDNA-seq data and initial SNV clustering *ψ*. This leads to the following problem statement.

#### Problem 1

(Clone Tree Selection (CTS)) Given a candidate set of clone trees 𝒯, an initial SNV clustering *ψ* : [*m*] → [*k*], and variant and total read count matrices 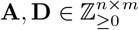 for *n* single cells and *m* SNVs, find the clone tree *T*^∗^ ∈ 𝒯 that maximizes the posterior probability *P* (*T* |**A, D**, *ψ*).

Using Bayes’ rule, and assuming a uniform prior over *T* ∈ 𝒯 and that the total read counts **D** are independent of *T* and *ψ*, i.e., *P* (**D** | *T, ψ*) = *P* (**D**), we have

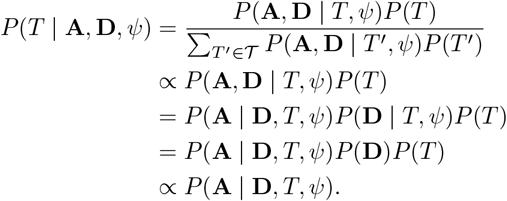

In our generative model, each cell *i* is generated from either a clone *q* in tree *T*, with *k* nodes or from an artificial root representing a normal cell not carrying any of the somatic mutations. Let *z*_*i*_ ∈ [*k* + 1] be the latent variable indicating the clone from which cell *i* was generated. Here, clone *k* + 1 represents the normal cells. Additionally, we assume our initial SNV clustering *ψ* is not error-free and that some SNVs may have been incorrectly clustered. Let *y*_*j*_ ∈ [*k*] be the latent variable indicating that SNV *j* belongs to some cluster *p* ∈ [*k*]. Since the cell-to-clone labels **z** and SNV-to-cluster labels **y** are unknown latent variables, we marginalize over the latent labels to compute the marginal likelihood

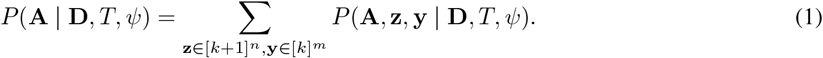

We model the observed variant read counts *a*_*i,j*_ using a binomial distribution. Given total reads *d*_*i,j*_, clone tree *T*, cell label *z*_*i*_ = *r*, and SNV cluster label *y*_*j*_ = *p*, the likelihood of observing variant reads *a*_*i,j*_ is

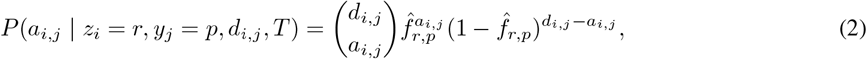

where the effective success probability 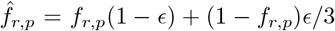 adjusts the latent variant allele frequency *f*_*r,p*_ of SNV cluster *p* in clone *r* for sequencing error rate *ϵ* [29, 30]. Since the latent variant allele frequency *f*_*r,p*_ for clone *r* and an SNV *j* in SNV cluster *p* is often unknown, we approximate the latent VAF *f*_*r,p*_ with 1*/*2 whenever SNV cluster *p* is present in clone *r* and 0 otherwise.

### 2.2 Method description

Direct evaluation of (1) requires summing over up to (*k* +1)^*n*^*k*^*m*^ latent configurations and is therefore computationally intractable. To solve the Clone Tree Selection (CTS) problem, we propose a variational Bayesian approach named Arborist. Our goal is to derive a tractable approximation to the marginal likelihood (1) for a fixed tree *T* ∈ 𝒯. To achieve a tractable evidence lower bound (ELBO) on the marginal likelihood, Arborist uses mean-field variational inference [36–38], which requires approximating the posterior *P* (**z, y** | **A, D**, *T, ψ*), obtained by applying Bayes’ rule to the joint likelihood in (1), with a factorized variational distribution *q*(**z, y**) [36–38].

Taking the logarithm of the marginal likelihood (1) and applying Jensen’s inequality yields a variational bound, i.e., the evidence lower bound (ELBO), on the marginal likelihood,

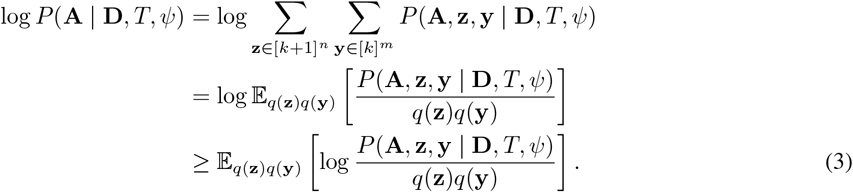

We parameterize the variational family using categorical distributions. Specifically, *q*_*ϕ*_(*z*_*i*_) = Categorical(***ϕ***_*i*_) and *q*_*θ*_(*y*_*j*_) = Categorical(***θ***_*j*_), where ***ϕ***_*i*_ and ***θ***_*j*_ are variational probability vectors for the latent cell-to-clone and SNV-to-cluster labels, respectively. To incorporate uncertainty in the initial SNV clustering *ψ*, we define a prior *π*_*j,p*_ = *P* (*y*_*j*_ = *p* | *ψ, γ*) as

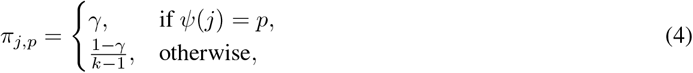

where *γ* ∈ [0, 1] reflects the confidence in the initial cluster assignment *ψ*. Alternatively, ***π***_*j*_ may be set directly from posterior cluster probabilities provided by bulk-based SNV clustering methods [15, 25, 35, 39].

Arborist seeks variational distributions *q*_*ϕ*_(**z**) and *q*_*θ*_(**y**) that maximize the ELBO denoted by ℒ (*q*_*ϕ*_, *q*_*θ*_). Under the mean-field factorization, the ELBO ℒ (*q*_*ϕ*_, *q*_*θ*_) can be written as

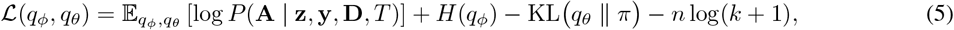

where *H*(*q*_*ϕ*_) is the entropy of the cell-to-clone posterior and KL(*q*_*θ*_ ∥ *π*) is the Kullback-Leibler divergence between the variational SNV-cluster distribution *q*_*θ*_ and the SNV-to-cluster prior ***π***.

The ELBO ℒ (*q*_*ϕ*_, *q*_*θ*_) decomposes over cells, SNVs, clones and SNV clusters under our factorization, such that

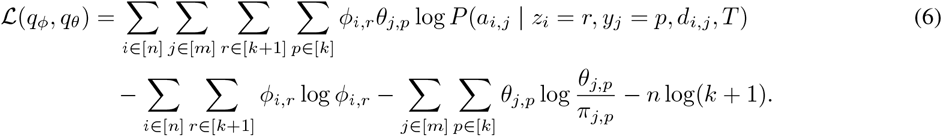

We optimize the ELBO (6) using coordinate ascent variational inference (CAVI) [36–38]. At iteration *t*, we alternately update 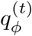 and 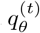 by coordinate updates of the variational parameters ***ϕ***^(*t*)^ and ***θ***^(*t*)^ as follows:

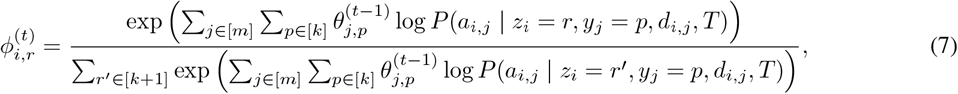

given ***θ***^(*t*−1)^ and

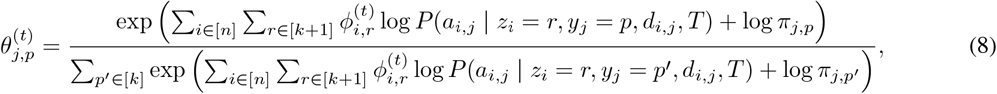

given ***ϕ***^(*t*)^. See Appendix A for the full derivation of the ELBO and the corresponding coordinate-ascent update equations. Let ℒ ^(*t*)^ denote the ELBO 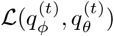 at iteration *t*. We repeat this process until convergence, i.e., | ℒ ^(*t*)^ – ℒ ^(*t*−1)^ | *< δ*, where *δ* ≥ 0 is a convergence tolerance, or until a maximum number of iterations. After convergence, we obtain point estimates of the latent assignments by taking the maximum a posteriori (MAP) assignments from the variational posteriors 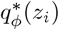 and 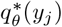.

A practical advantage of variational inference is that the ELBO naturally supports model selection across clone trees of different sizes [37,38,40]. Because the ELBO is a lower bound on the marginal likelihood and includes explicit complexity penalties through the entropy and KL divergence terms, it balances goodness-of-fit against unnecessary latent structure without requiring any additional regularization. This makes the ELBO a principled way to compare heterogeneous candidate sets produced by bulk methods particularly in settings where inference procedures, such as Conipher, may add or remove SNV clusters, producing trees of varying sizes.

This variational inference procedure is performed independently for each candidate tree *T*∈ 𝒯, with Arborist returning the clone tree *T*^∗^ with the maximum ELBO and the corresponding MAP SNV-to-cluster labels **y** and cell-to-clone labels **z**. Arborist is implemented in Python, is open source, and available at https://github.com/VanLoo-lab/Arborist.

## 3 Results

### 3.1 Benchmarking

We developed Arborist, which makes use of variational inference to solve the Clone Tree Selection problem. To benchmark Arborist against existing bulk sequencing and scDNA-seq only clone tree inference methods, we performed a simulation study under various sequencing regimes and characteristics of a tumor. First, we generated a ground truth clone tree *T* as described in Weber *et al*. [30] varying the number of clones *k* ∈ {7, 10}. Each node *u* ∈ *V* (*T*) and SNV *j* ∈ [*m*] has an associated genotype *g*(*u, j*) = (*x, y, w*) for clone *u* that captures both the allele specific copy number (*x, y*) ∈ (Z_*≥*0_, Z_*≥*0_) and the number of mutated copies *w* ∈ {0, …, max(*x, y*)} for each of the *m* = 10000 SNVs. Given a simulated clone tree *T*, we sampled clonal proportions according to a Dirichlet distribution for *p* = 3 bulk samples and then simulated bulk read counts with coverage ∈ {50×, 100×}. Lastly, we simulated single-cell read counts with coverage ∈ {0.02×, 0.05×} for *n* = 1500 cells, with 500 cells per sample. Although Ar-borist utilizes the infinite sites assumption, the simulated clone trees contain SNV loss via copy number aberrations for approximately 10% of the *m* SNVs.

To obtain an initial SNV clustering, we utilized PyClone-VI [15, 39] due to its widespread adoption within the community. For bulk clone tree inference methods, we benchmarked against Conipher [20], due to its use in analysis of the TRACERx cohort [4, 14], and Sapling [24, 41], which is able to return a large candidate set. In particular, we utilized these bulk inference methods in three ways. First, we sought to evaluate how pairing Arborist with either Conipher or Sapling could improve the accuracy of clone tree reconstruction as well as the accuracy of the cell-to-clone labels and SNV-to-cluster labels. These approaches are referred to as Conipher+Arborist and Sapling+Arborist, respectively.

For all experiments, we used a per-base sequencing error rate of 0.001 and set the SNV-to-cluster prior ***π***_*j*_ (4) with *γ* = 0.9. We also used the default Arborist convergence tolerance of *δ* = 1. However, since Conipher may remove SNV clusters during tree inference, we instead used a uniform SNV-to-cluster prior ***π***_*j*_ for each SNV *j* in the Conipher removed clusters. This ensured that Arborist still approximated the SNV-to-cluster posterior distribution for all SNVs. Second, we utilized the default tree in the case of Conipher or the highest likelihood tree returned by Sapling. These methods are referred to as Conipher and Sapling. Lastly, we utilized the solution space of trees returned by Conipher and Sapling in order to find a single consensus tree using GraPhyC [26]. Since Conipher, Sapling and GraPhyC do not provide cell-to-clone labels for single cells, we computed the likelihood of each cell originating from each clone in the Conipher or Sapling clone trees and assigned each cell to the clone with the highest likelihood in order to compare against Arborist. For scDNA-seq only tree inference methods, we benchmarked against Phertilizer [29] and SBMClone [31]. Since Phertilizer requires copy number profiles or binned read counts, we provided Phertilizer with the ground truth simulated copy number profiles of each cell, giving it an inherent advantage. Since SBMClone does not infer a clone tree, we utilized a similar procedure to that described in McPherson *et al*. [10] to convert the SBMClone block matrix to a clone tree with minor modifications to automate any manual steps—see Appendix B for more details.

To evaluate clone tree reconstruction accuracy, we computed three recall-based metrics that capture pairwise SNV (cell) relationships. Specifically, we measured (i) the *ancestral pair recall* (APR), defined as the fraction of ground-truth ordered SNV pairs that are in an ancestral relationship in the inferred tree *T* ^*′*^, (ii) the *incomparable pair recall* (IPR), defined as the fraction of ground-truth unordered SNV pairs that occur on different branches, i.e., are incomparable in *T* that are also incomparable in the inferred tree *T* ^*′*^, and (iii) the *clustered pair recall* (CPR), defined as the fraction of ground-truth unordered SNV pairs that belong to the same cluster in ground-truth tree *T* that are also clustered together in the inferred tree *T* ^*′*^. Additionally, we computed the adjusted rand index (ARI) on the SNV-to-cluster and cell-to-clone labels with respect to the ground truth assignments.

We focus our discussion on benchmarking with *k* = 10 clusters, a bulk sequencing depth of 50× and scDNA-seq coverage of 0.02× as this is the most challenging regime tested. When benchmarking against bulk sequencing only clone tree inference methods, we observed that the inclusion of Arborist, i.e., Conipher+Arborist and Sapling+Arborist, boosted performance over their counterparts Conipher and Sapling (Fig. 2). Additionally, both Conipher+Arborist and Sapling+Arborist outperformed the consensus method GraPhyC on all performance metrics. The greatest improvement in performance was on the SNV IPR metric (Fig. 2b) with Conipher+Arborist (median:0.90) and Sapling+Arborist (median: 0.91) outperforming their counterparts, Conipher (median: 0.71) and Sapling (median: 0.70). GraPhyC (median: 0.77) slightly outperformed both Conipher and Sapling on the IPR metric. We observed similar trends for the cell IPR metric (Fig. 2b) with Conipher+Arborist (median: 0.89) and Sapling+Arborist (median: 0.91) again having the top performance over Conipher (median: 0.43), Sapling (median: 0.58) and GraPhyC (median: 0.57). On the SNV (cell) APR and SNV (cell) CPR metrics, we found that the inclusion of Arborist led to improved performance over Conipher, Sapling and GraPhyC. Notably, Arborist was able to significantly improve the initial Pyclone-VI clustering (median: 0.55) as evidenced by SNV ARI for Conipher+Arborist (median: 0.75) and Sapling+Arborist (median: 0.73). One explanation for why Conipher+Arborist slightly outperforms Sapling+Arborist is that Conipher removes SNV clusters during tree inference while Sapling does not and ultimately infers clone trees much closer to the ground truth number of SNV clusters (Fig. S1). The SNVs in the SNV clusters excluded by Conipher are then redistributed by Arborist to a better fitting existing SNV cluster and PyClone-VI errors are less likely to be propagated to the clone tree inference step. Arborist also outperformed both scDNA-seq only inference methods, Phertilizer and SBMClone. The difference in performance was especially apparent in the lowest coverage of 0.02×, which is representative of the typically sequencing coverage of current scDNA-seq datasets. While SBMClone (median: 0.76) performed comparably on SNV clustering to Conipher+Arborist (0.76) and SNV CPR (Conipher+Arborist median: 0.82 vs. SBMClone median: 0.84), SBMClone (median APR: 0.68, median IPR: 0.60) was not competitive with Conipher+Arborist (median APR: 0.92, median IPR: 0.90) and Sapling+Arborist (median APR: 0.92, median IPR: 0.91) in terms of correctly identifying the ancestral and incomparable relationship of SNVs. Similarly, Phertilizer had top performance on SNV CPR (median: 0.87) and cell CPR (median: 0.88) due to its tendency to underestimate the number of SNV clusters/clones (Fig. S1), but otherwise was not competitive with the top performing methods on all other metrics.

**Figure 2.**
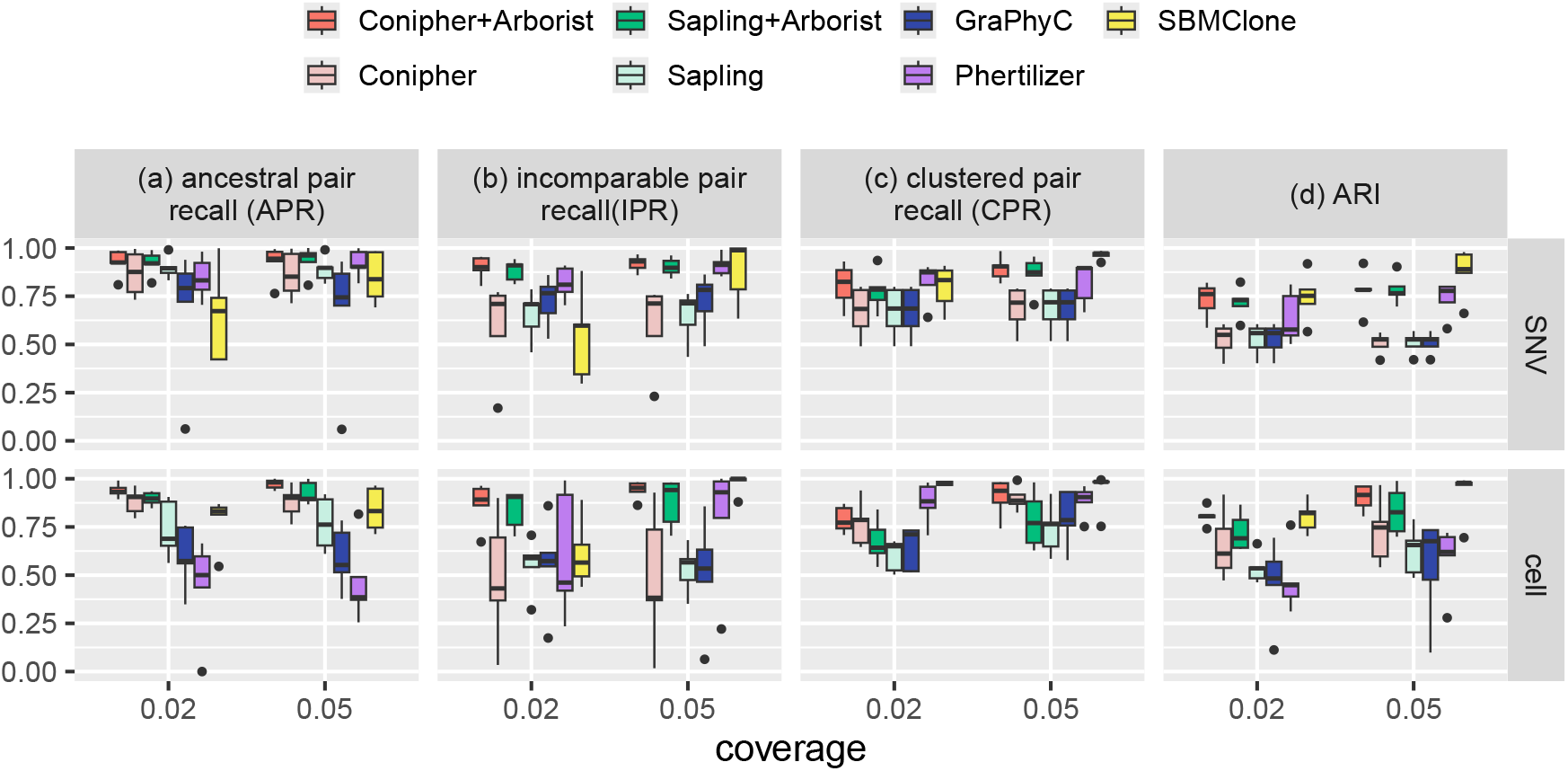
Arborist improves accuracy of clone tree reconstruction on simulated data. Results shown for *k* = 10 SNV clusters and bulk sequencing coverage of 50 ×. PyClone-VI [15,39] was used to find the initial SNV clustering *ψ* for all bulk sequencing methods. (a) SNV and cell ancestral pair recall, (b) SNV and cell incomparable pair recall, (c) SNV and cell clustered pair recall and (d) SNV-to-cluster label and cell-to-clone label adjusted Rand index (ARI).

We observed similar trends for *k* = 7 SNV clusters, and/or bulk sequencing coverage of 50 × /100× —see Figs. S2-S7. We also note that while the relative performance of scDNA-seq-only methods improve as scDNA-seq sequencing coverage increases from 0.02× to 0.05×, both Phertilizer and SBMClone are unable to outperform the combination of PyClone-VI, Conipher and Arborist on clone tree reconstruction metrics. Overall, the inclusion of Arborist improved the accuracy of the clone tree reconstruction as well as the accuracy of cell-to-clone labels and SNV-to-cluster labels over bulk sequencing and scDNA-seq-only methods, especially in the ultra-low coverage (0.02×) regime. This demonstrates the utility of jointly using bulk sequencing and scDNA-seq data in a principled way to obtain more accurate clone trees for downstream evolutionary analysis.

### 3.2 Application of ARBORIST to a Malignant Peripheral Nerve Sheath Tumor

To validate Arborist on biological data, we analyzed a malignant peripheral nerve sheath tumor (MPNST), a rare sarcoma originating from Schwann cells [11,42]. We performed multi-region sequencing for 5 spatially separated samples of patient GEM2.3 and called 8563 SNVs using two-out-of-three consensus calling via Mutect2 [43], Strelka [44] and MuSE [45]. We clustered SNVs using DPClust [35] and then utilized the union of clone trees inferred by both Conipher and Sapling to obtain a candidate set 𝒯 of 100 clone trees. We also sequenced 3190 single cells with ACT [8] sampled from the 5 distinct regions. Of the 8563 SNVs called from the bulk sequencing data, only 4767 SNVs had at least one mapped alternate read within the scDNA-seq. We then used Arborist to rank the candidate set and obtain MAP assignments for the cell-to-clone labels and SNV-to-cluster labels. A *γ* value of 0.9 was used to set the SNV-to-cluster prior to indicate high confidence in the initial SNV clustering *ψ* since the DPClust results had been manually validated. The results are shown in Fig. 3.

**Figure 3:**
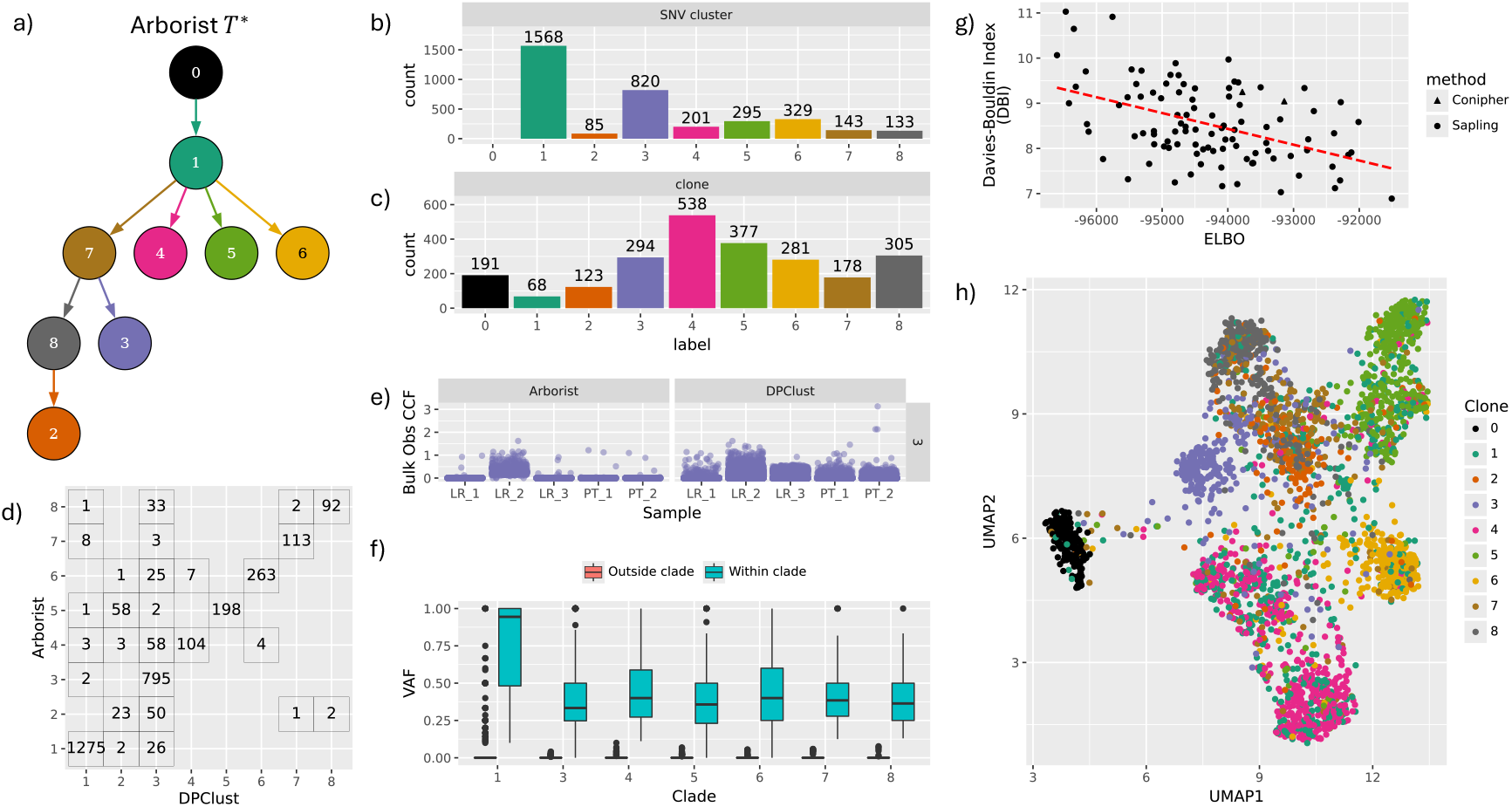
Arborist results for MPNST patient GEM2.3. (a) The clone tree *T*^∗^ with maximum ELBO selected by Arborist. Nodes and incoming edges are colored by clone and SNV cluster label, respectively. Note that label 0 represents normal cells. (b) The distribution of SNV-to-cluster labels **y** corresponding to clone tree *T*^∗^. (c) The distribution of cell-to-clone labels **z** corresponding to clone tree *T*^∗^. (d) The comparison between DPClust initial cluster labels and Arborist SNV-to-cluster labels **y**^∗^. (e) The distribution of bulk observed CCFs by sample for SNV cluster 3 grouped by clustering method. (f) The distribution of *within clade* and *outside clade* VAFs by clade in clone tree *T*^∗^. (g) The relationship between the ELBO and the Davies-Bouldin Index computed for normalized binned read counts and MAP cell-to-clone labels **z** computed for each clone tree *T*∈ 𝒯. (h) UMAP of normalized binned read counts with points colored by cell-to-clone label **z**.

The top ranked tree returned by Arborist originated from Sapling and is shown in Fig. 3a. Interestingly, of the two clone trees generated by Conipher, the first tree returned by Conipher had a worse Arborist rank (28) when compared to the second tree (16). For analysis, we exclude (i) cells in the lowest 10th percentile of total mapped reads, and (ii) cells and SNVs whose variational entropy, defined as 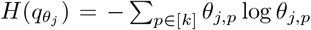 for SNVs and similarly 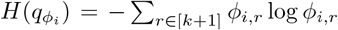 for cells, lies in the top 75th percentile (Figs. S8, S9). Fig. 3b,c show the distribution of the MAP SNV-to-cluster labels **y** and cell-to-clone labels **z**. The clonal cluster, SNV cluster 1, had the largest number of SNVs (1568), while SNV cluster 2 was the smallest with only 85 confidently placed SNVs. For clones, the largest clone was clone 4 with 538 cells while clone 1 had the fewest number of cells (68).

Next, we compared the Arborist SNV-to-cluster labels **y** to the initial DPClust labels (Fig. 3d). Overall, Ar-borist retained the structure of the initial clustering with the exception of SNV cluster 3, where Arborist refines SNV cluster 3 by redistributing SNVs to existing clusters. We compared the distribution of the cancer cell fraction (CCF) [46] observed in each of the 5 bulk samples by SNV cluster (Fig. 3e, S10), demonstrating a reduction of noise in the distribution of observed CCF by sample and indicating that SNV cluster 3 is private to sample LR-2 as opposed to all samples.

To assess the goodness of fit of Arborist selected clone tree *T*^∗^, we computed a *within clade VAF v*_*u,j*_ and an *outside clade* 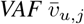 for each clade *u* and SNV *j*, defined respectively as

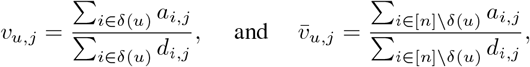

where set *δ*(*u*) contains cells mapped to clones in clade *u*. Intuitively, we expect *within clade VAF* to be high since SNV *j* should be present within this set of cells and the *outside clade VAF* to be near zero since the SNVs should not be present in these cells. We observed a clear separation between *within clade VAF* and *outside clade VAF* for each clade in clone tree *T*^∗^ (Fig. 3f), providing further supporting evidence for correct assignment of SNVs and cells within clone tree *T*^∗^.

Lastly, because Arborist does not incorporate copy number information during inference, we orthogonally validated the resulting clone tree rankings using a proxy for copy number. Since low-pass scDNA-seq has uniform sequencing coverage, binned read counts serves a reasonable proxy for the copy number profile of a single cell. For each cell and each 500 kb genomic bin within the autosomes, we counted the number of reads mapping to that bin and normalized by the total number of reads per cell. We then removed bins in the 95th percentile of variance across cells to reduce noise. For each cell-to-clone labeling **z** returned by Arborist, we computed the Davies–Bouldin Index (DBI) [47], a standard clustering metric that evaluates the average similarity of each cluster to its most similar other cluster, where similarity is defined as the ratio of within-cluster dispersion to between-cluster separation. Lower DBI values correspond to better defined clusters, implying low within-clone variability and high separation between clones.

As shown in Fig. 3g, higher ELBO values correspond to lower DBI, indicating that better ranked Arborist clone trees also exhibit more coherent clustering patterns in normalized binned read count space. Notably, the Arborist selected clone tree *T*^∗^ achieved the lowest overall DBI of 6.9. To visually assess cluster coherence, we performed UMAP (Fig. 3h) on the normalized binned read counts and colored cells by the Arborist cell-to-clone labels **z**^∗^. Despite normalized binned read counts being a relatively noisy proxy for copy number, clear patterns of clonal structure and separation between clusters remain readily observable in the UMAP embedding.

Overall, we found strong evidence for the Arborist selected clone tree *T*^∗^ and its corresponding SNV-to-cluster labels **y**^∗^ and cell-to-clone label **z**^∗^ via validation analysis of both SNVs and an orthogonal proxy for copy number. Additionally, since SNV clustering on bulk sequencing remains an ongoing challenge in the field due to the need for deconvolution of clonal populations, Arborist effectively denoised the original DPClust SNV clusters, yielding improved SNV clusters for downstream analysis.

## 4 Discussion

In this work, we introduced the Clone Tree Selection problem, which formalizes the challenge of resolving ambiguity within the clone tree solution space inferred from bulk data alone. To solve the CTS problem, we proposed Arborist, which computes a variational lower bound on the marginal likelihood of the scDNA-seq data given a fixed clone tree *T* and initial clustering *ψ*, and uses this bound to rank candidate clone trees and obtain corresponding MAP SNV-to-cluster and cell-to-clone labels. We demonstrated on simulated data that including Arborist in the clone tree inference pipeline, whenever scDNA-seq data is available, yields a more accurate reconstruction of the evolutionary history of the tumor than via bulk or single-cell inference methods alone. We also showed the utility of Arborist on a patient with MPNST, using it to discriminate between a large candidate set of clone trees and further refining the SNV clustering. With the increasing accessibility of low-pass scDNA-seq, Arborist provides a principled and scalable framework for integrated bulk and scDNA-seq clone tree inference, serving as a practical alternative to fully joint probabilistic modeling approaches, while maintaining accuracy for downstream evolutionary analyses.

There are a number of directions for future work. First, while we demonstrated the use of Arborist for joint bulk sequencing and scDNA-seq data, the framework could also be used as an scDNA-seq-only inference method when bulk sequencing data are not available. In this setting, single cells could be pooled into a pseudobulk sample, existing bulk sequencing analysis methods applied, and Arborist then used to select the best clone tree and improve the SNV clustering. Although pseudobulk samples often have lower effective depth and thus noisier SNV calls and clusters, Arborist in principle may help denoise and stabilize pseudobulk-based inference. Second, currently Arborist assumes a fixed clone tree, however an additional tree refinement step, similar to that used by Phertilizer [29], could enable more precise inference of SNV clusters and clones directly from scDNA-seq data. Third, the Arborist generative model could be extended to incorporate variant and total read counts from bulk sequencing data, as well as variational distributions to approximate SNV cluster CCF posterior distributions. Fourth, accuracy could be further improved by integrating copy number or binned read count information, allowing Arborist to prioritize clone trees whose inferred clones exhibit coherent structure across both SNV and copy number features. Fifth, more complex priors on the initial SNV clustering could be explored, providing a more natural way to encode confidence in the input SNV clustering. Lastly, Arborist could be extended beyond the infinite-sites assumption to capture SNV loss and other complex mutational processes and events.

## 5 Author contributions

LLW and PVL conceived of the project. PVL supervised the project. LLW and CYC implemented the method. LLW, CYC, CL and CG conducted the experiments. LLW, YP and YC validated the results. LLW and PVL wrote and edited the manuscript. All authors read and approved the final manuscript.

## 6 Acknowledgments

PVL is a CPRIT Scholar in Cancer Research and acknowledges CPRIT grant support (RR210006).

## A Supplemental methods

### A.1 Derivation of the evidence lower bounds (ELBO)

We derive the evidence lower bound (ELBO) used by Arborist. Recall the marginal likelihood (1)

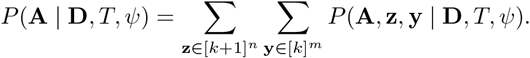

Let *q*(**z, y**) be any variational distribution over the latent variables. Applying Jensen’s inequality,

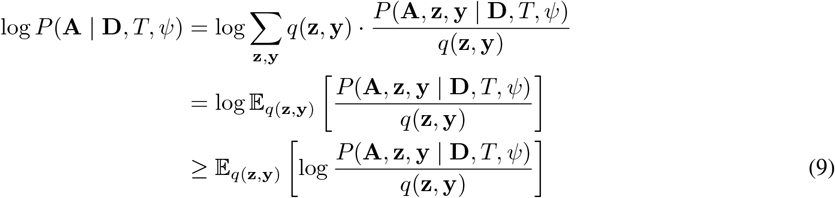

Thus, giving the evidence lower bound (ELBO)

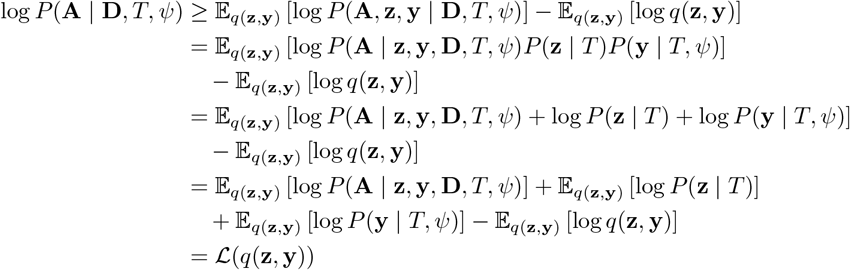

We use the mean-field factorization

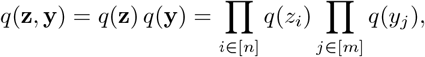

with categorical variational distributions

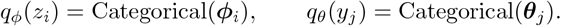

to rewrite the ELBO as

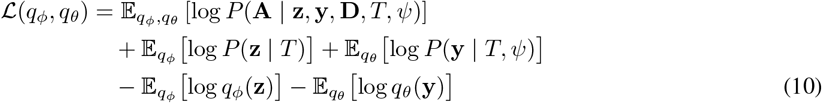

Noting the definition of KL-divergence,

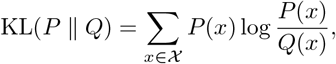

we can rearrange the expression above as

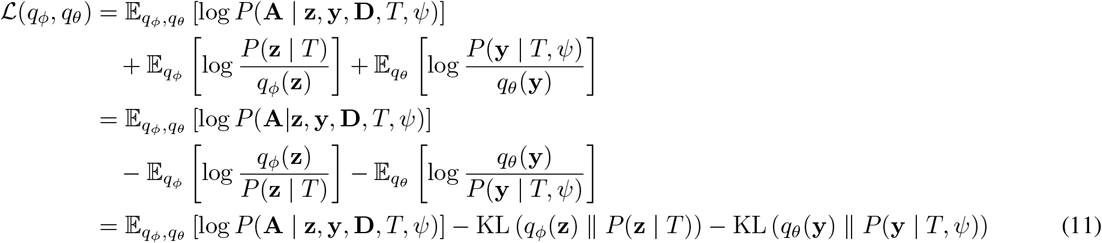

Assuming, conditional independence between cells and SNVs, we rewrite

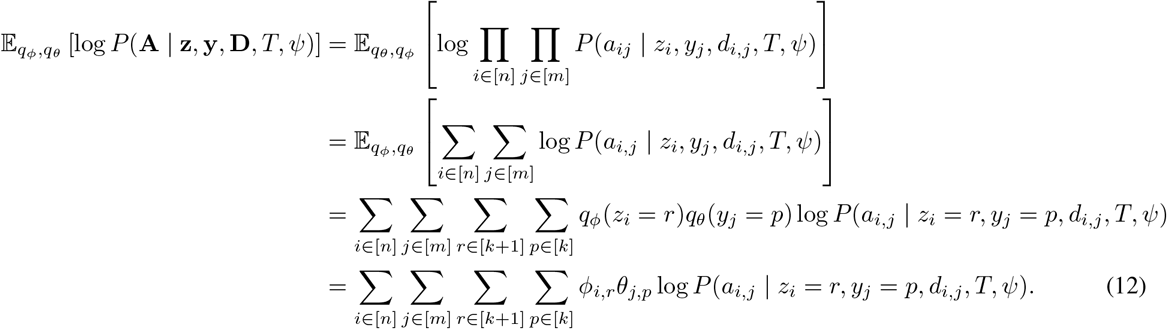

We assume a uniform prior over the *k* tumor clones and an additional normal clone for a total of *k* + 1 clone labels, such that

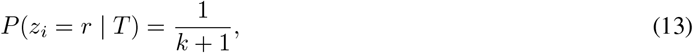

and then the KL divergence term is written as

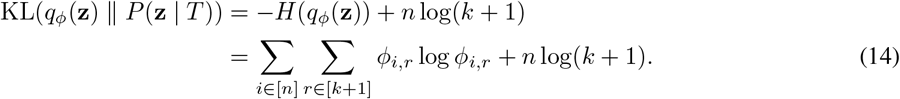

We define a prior on *P* (*y*_*j*_ = *p* |*T, ψ*) = *π*_*j,p*_ parameterized by *γ* ∈ [0, 1], which represents the confidence-level in the initial SNV clustering *ψ*. Thus,

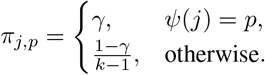

Thus, giving

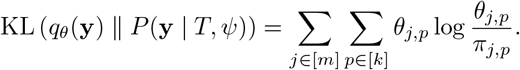

Putting together (12), (14) and (15), yields ELBO

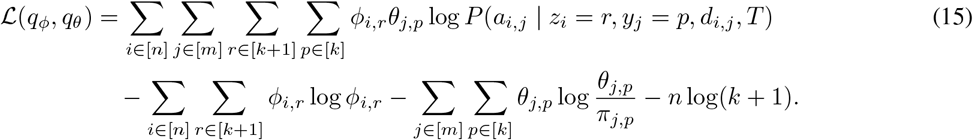

### A.2 CAVI update equations for variational distributions

Next, we derive the CAVI update for *q*_*θ*_(**y**) and *q*_*ϕ*_(**z**). Note that each variation distribution *q*_*θ*_(*y*_*j*_) and *q*_*ϕ*_(*z*_*i*_) must satisfy the normalization constraints, Σ_*p*∈[*k*]_ *θ*_*j,p*_ = 1 for each SNV *j* and Σ_*r*∈[*k*+1]_ *ϕ*_*i,r*_ = 1. To meet these constraints, we introduce Lagrange multipliers *λ*_*j*_ for each SNV *j* ∈ [*m*] and similarly *λ*_*i*_ for each cell *i* ∈ [*n*] and form the constrained Lagrangian

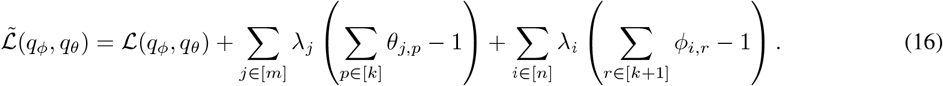

Taking the partial derivative with respect to *θ*_*j,p*_ yields

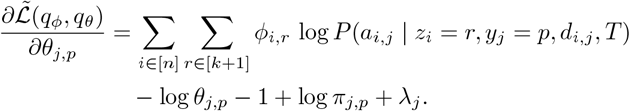

Setting the partial derivative to 0 and substituting in

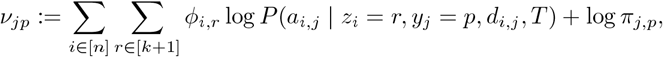

we obtain

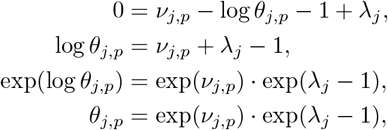

Then letting *C*_*j*_ = exp(*λ*_*j*_ − 1) and using that fact that Σ_*p*∈[*k*]_ *θ*_*j,p*_ = 1,

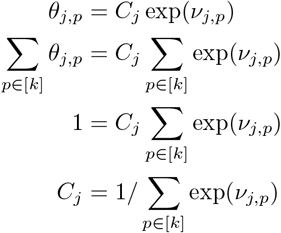

Substituting back yields the update equation for *θ*_*j,p*_,

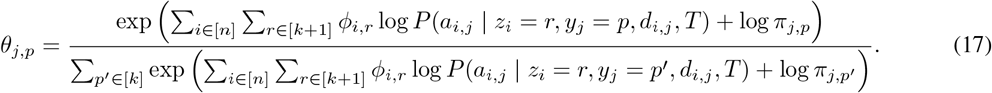

We now repeat these steps to derive the update equations for *q*_*ϕ*_(**z**) given *q*_*θ*_(**y**).

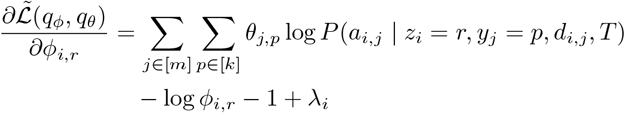

Setting *η*_*i,r*_ := Σ_*j* ∈ [*m*]_ Σ_*p*∈ [*k*]_ *θ*_*j,p*_ log *P* (*a*_*i,j*_ | *z*_*i*_ = *r, y*_*j*_ = *p, d*_*i,j*_, *T*) and setting the partial derivative to 0 yields

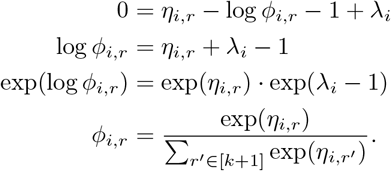

Finally, substituting back in for *η*_*i,r*_ yields the update equation for *ϕ*_*i,r*_ as

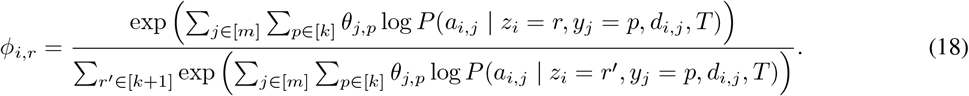

## B Benchmarking details

### B.1 Converting an SBMCLONE block matrix into a clone tree

We followed a similar procedure as McPherson *et al*.. [10] to convert the SBMClone block matrix into a clone tree under the infinite-sites assumption. We first binarized the block matrix by thresholding block probabilities at 0.01. We then removed all-zero rows and columns and merged duplicate rows and duplicate columns to obtain a reduced binary matrix. Next, we attempted to construct a perfect phylogeny on this matrix using the algorithm of Gusfield [48].

In McPherson *et al*., [10], if a block matrix violated the perfect phylogeny constraints, it was manually modified until it yielded a perfect phylogeny. Since manual correction was impractical for a large number of simulations, we instead developed a heuristic to modify the block matrix until it yielded a perfect phylogeny. First, we identified a conflicting 1-entry and flipped it to 0, choosing the entry with the lowest block probability. Then, this conflictresolution procedure was repeated until the modified matrix admitted a perfect phylogeny, which was then used as the inferred clone tree for benchmarking.

## C Supplemental figures

**Figure S1:**
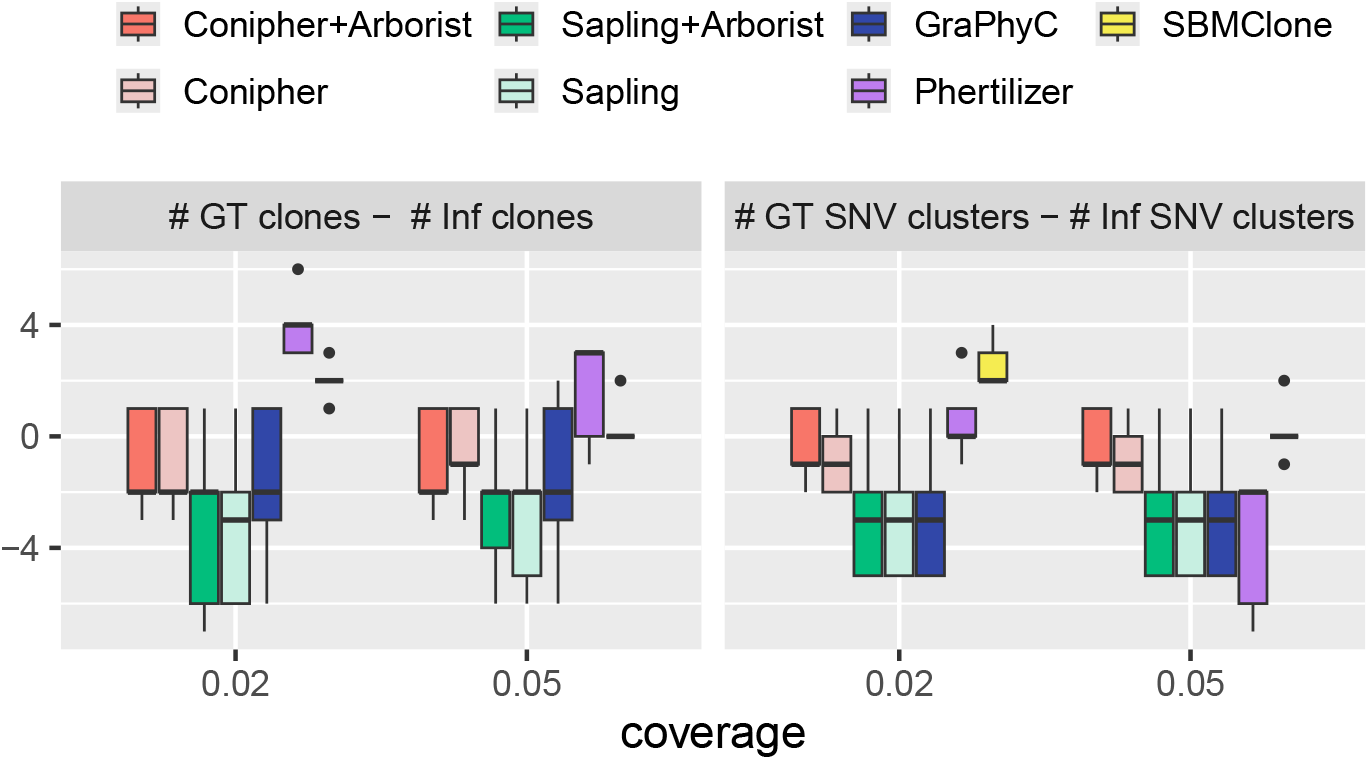
Difference in inferred non-empty clones and SNV clusters from ground truth for *k* = 10 SNV clusters and bulk sequencing depth of 50×.

**Figure S2:**
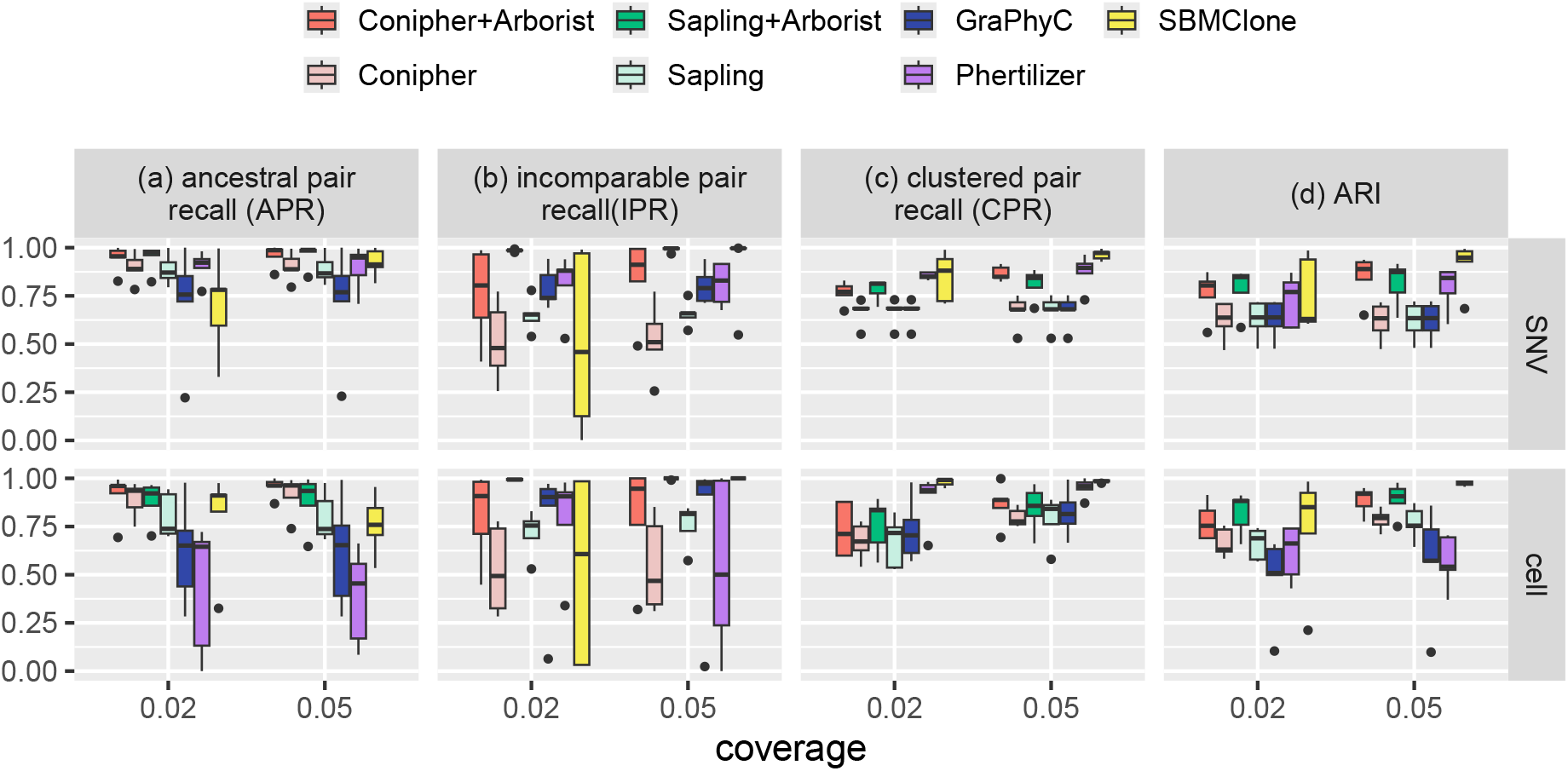
Benchmarking performance metrics for *k* = 7 SNV clusters and bulk sequencing depth of 50×.

**Figure S3:**
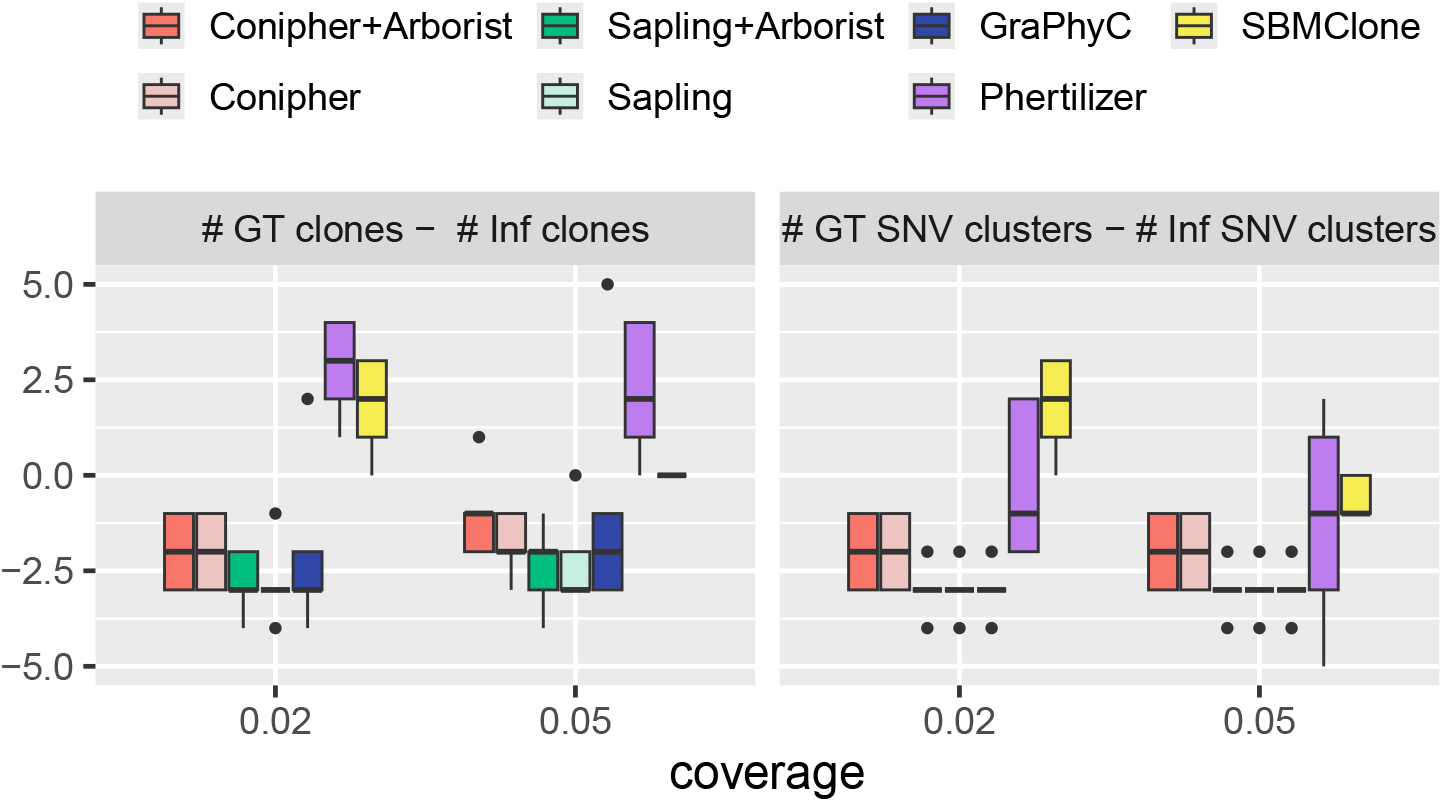
Difference in inferred non-empty clones and SNV clusters from ground truth for *k* = 7 SNV clusters and bulk sequencing depth of 50×.

**Figure S4:**
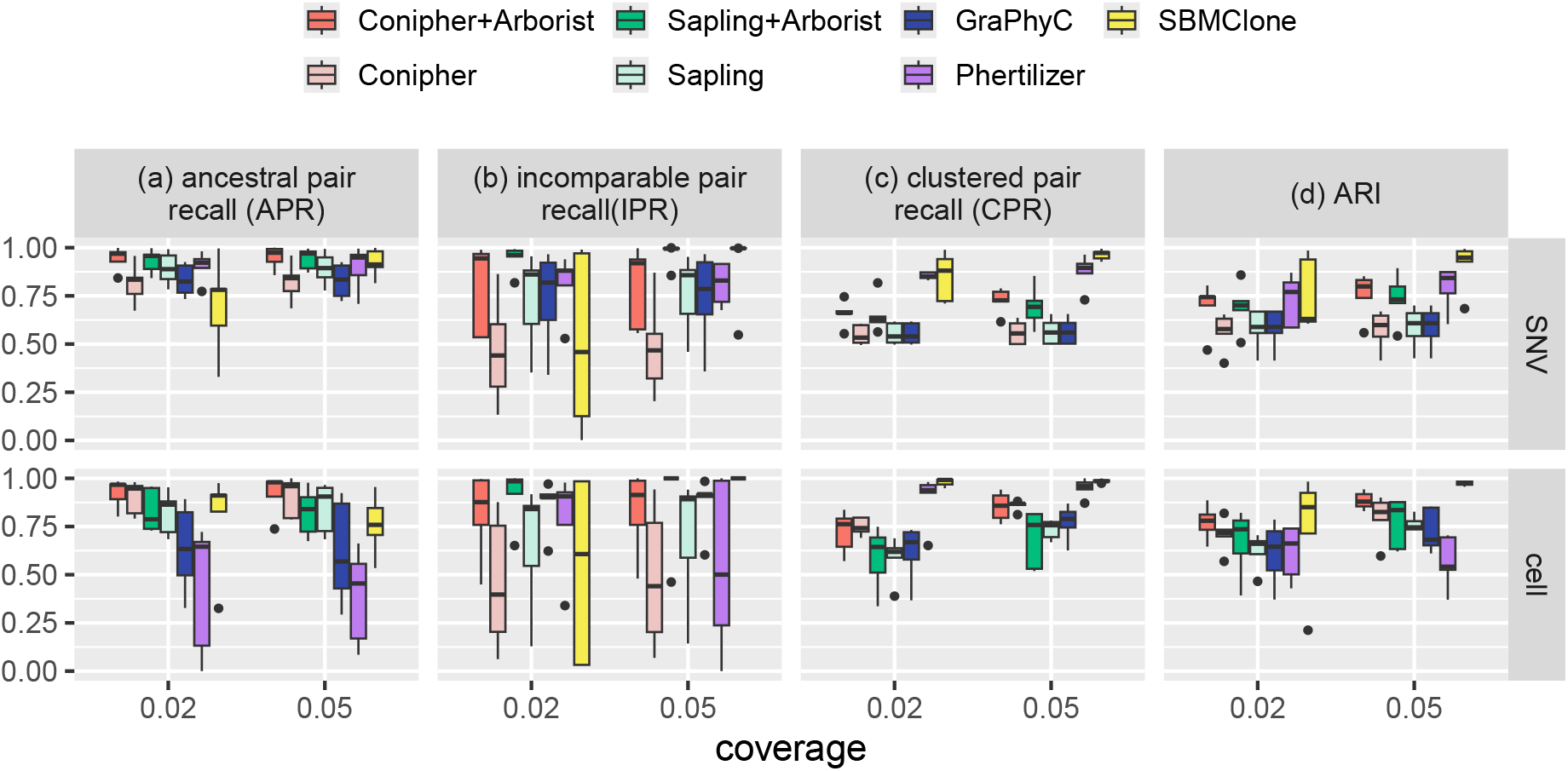
Benchmarking performance metrics for *k* = 7 SNV clusters and bulk sequencing depth of 100×.

**Figure S5:**
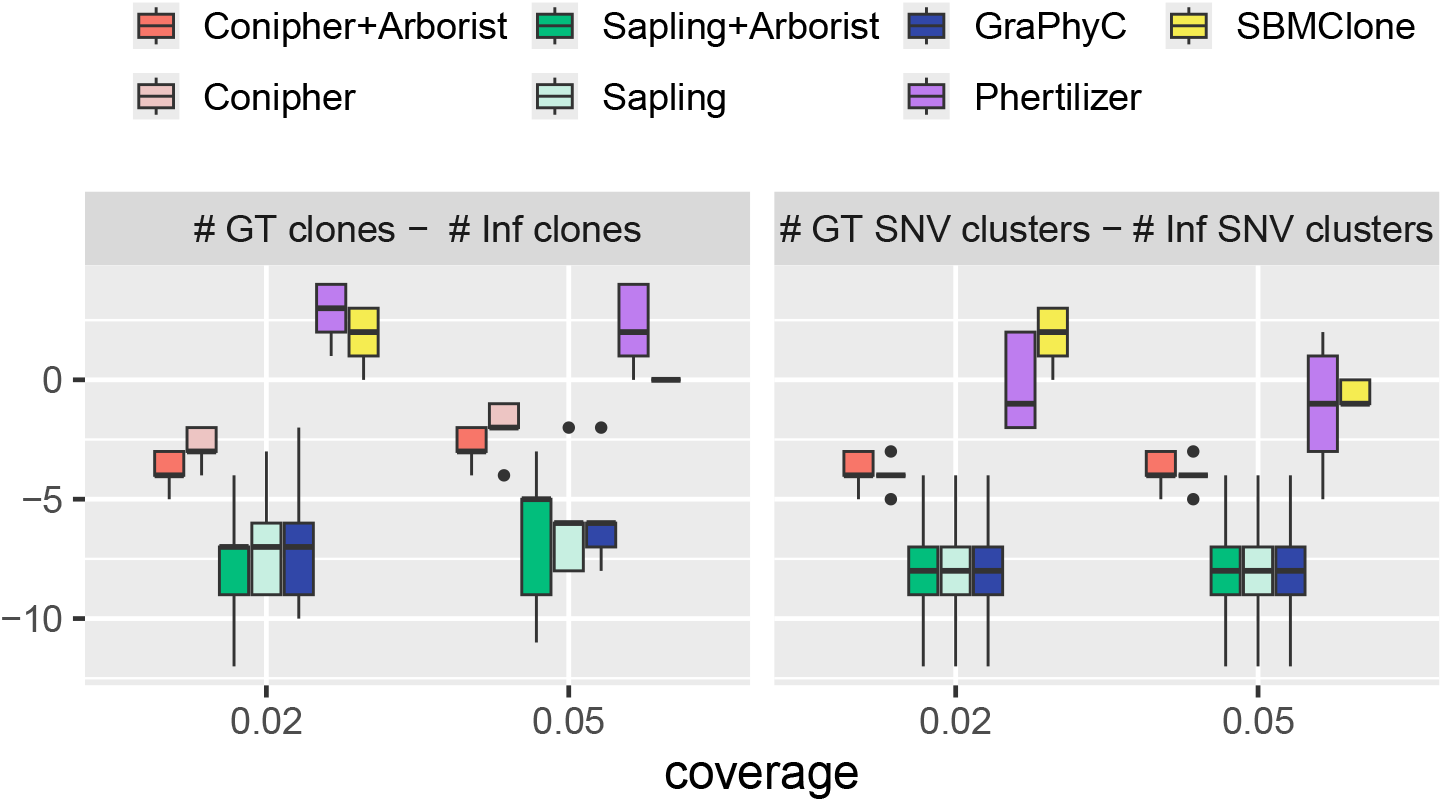
Difference in inferred non-empty clones and SNV clusters from ground truth for *.k* = 7 SNV clusters and bulk sequencing depth of 100×.

**Figure S6:**
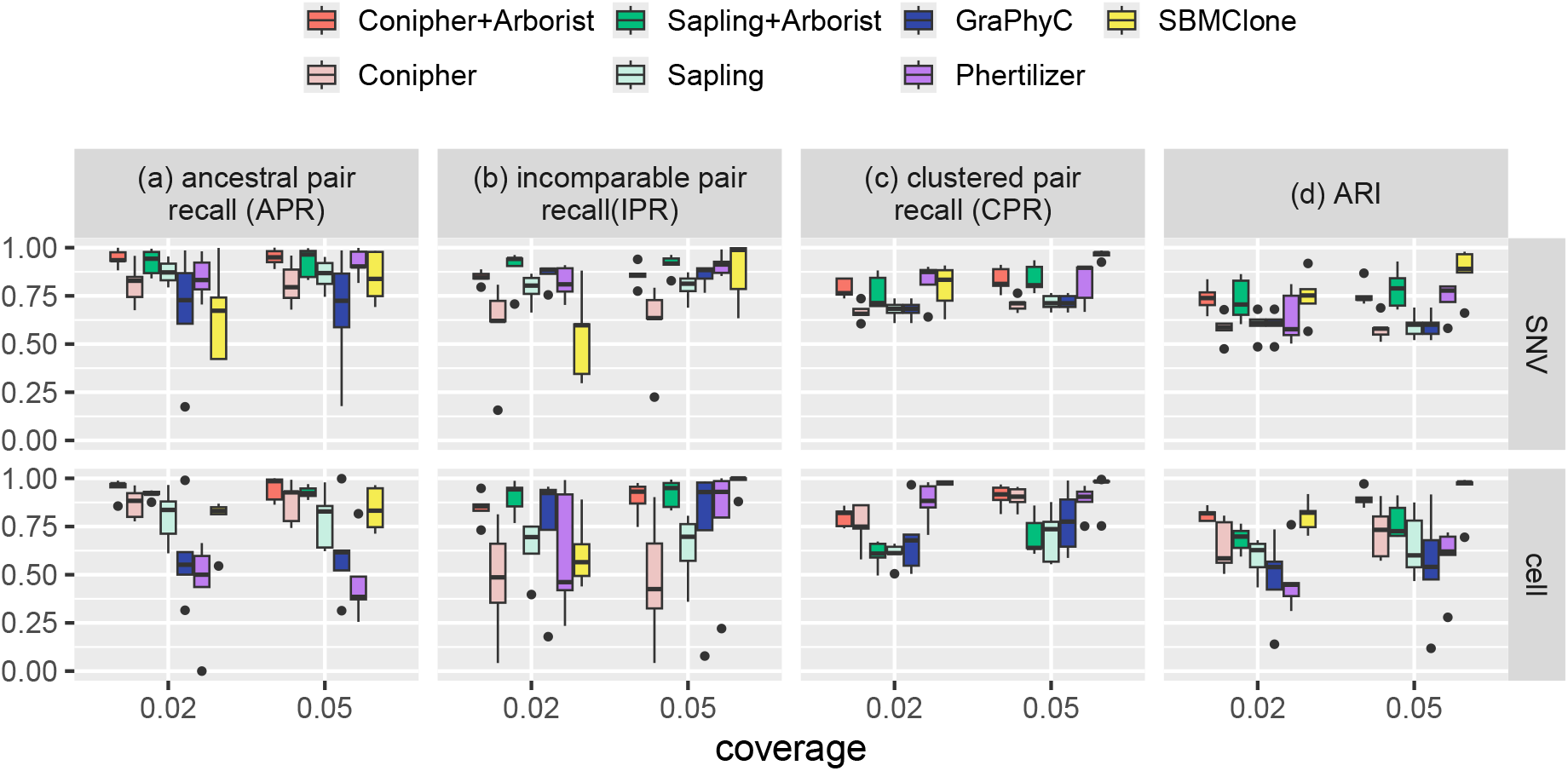
Benchmarking performance metrics for *k* = 10 SNV clusters and bulk sequencing depth of 100×.

**Figure S7:**
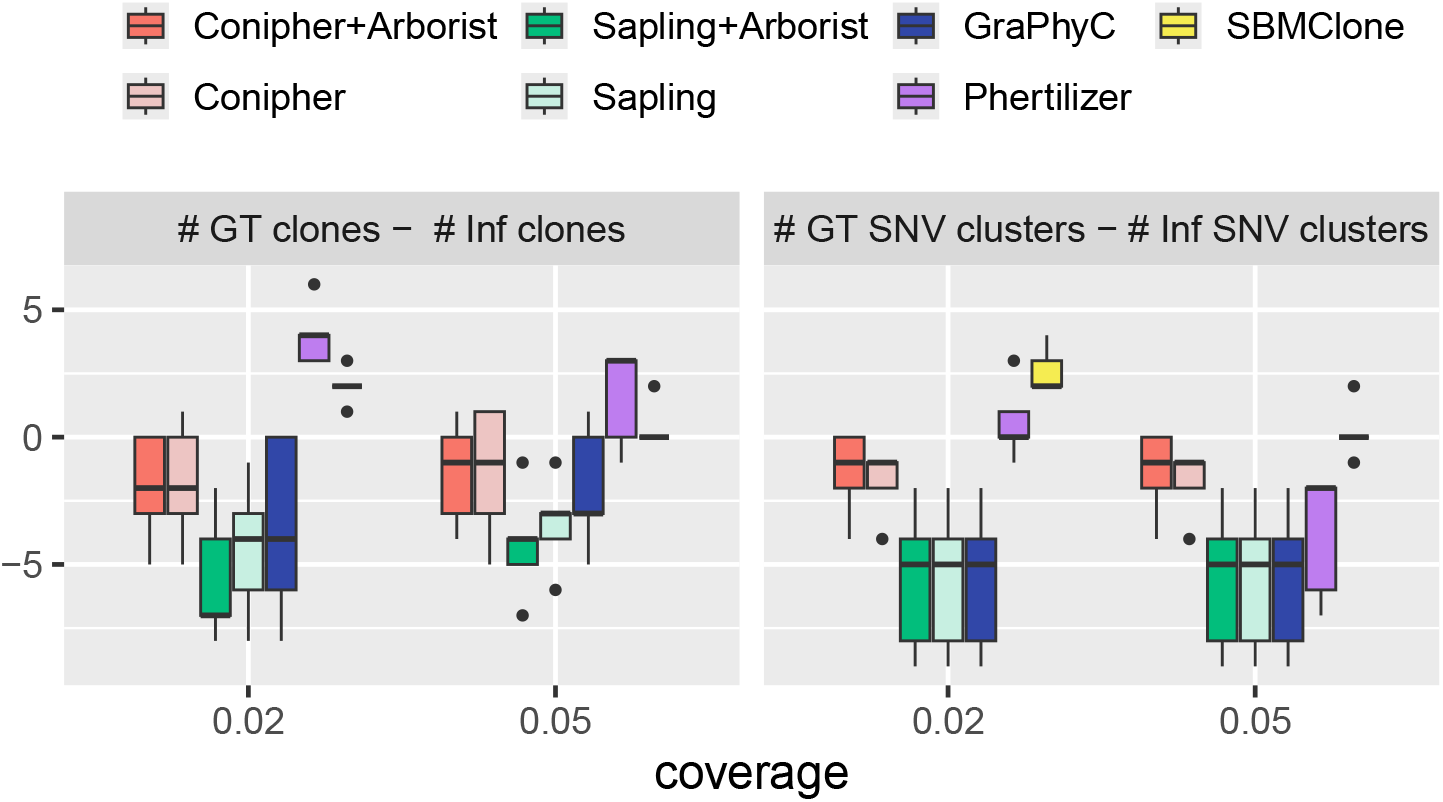
Difference in inferred non-empty clones and SNV clusters from ground truth for *k* = 10 SNV clusters and bulk sequencing depth of 100×.

**Figure S8:**
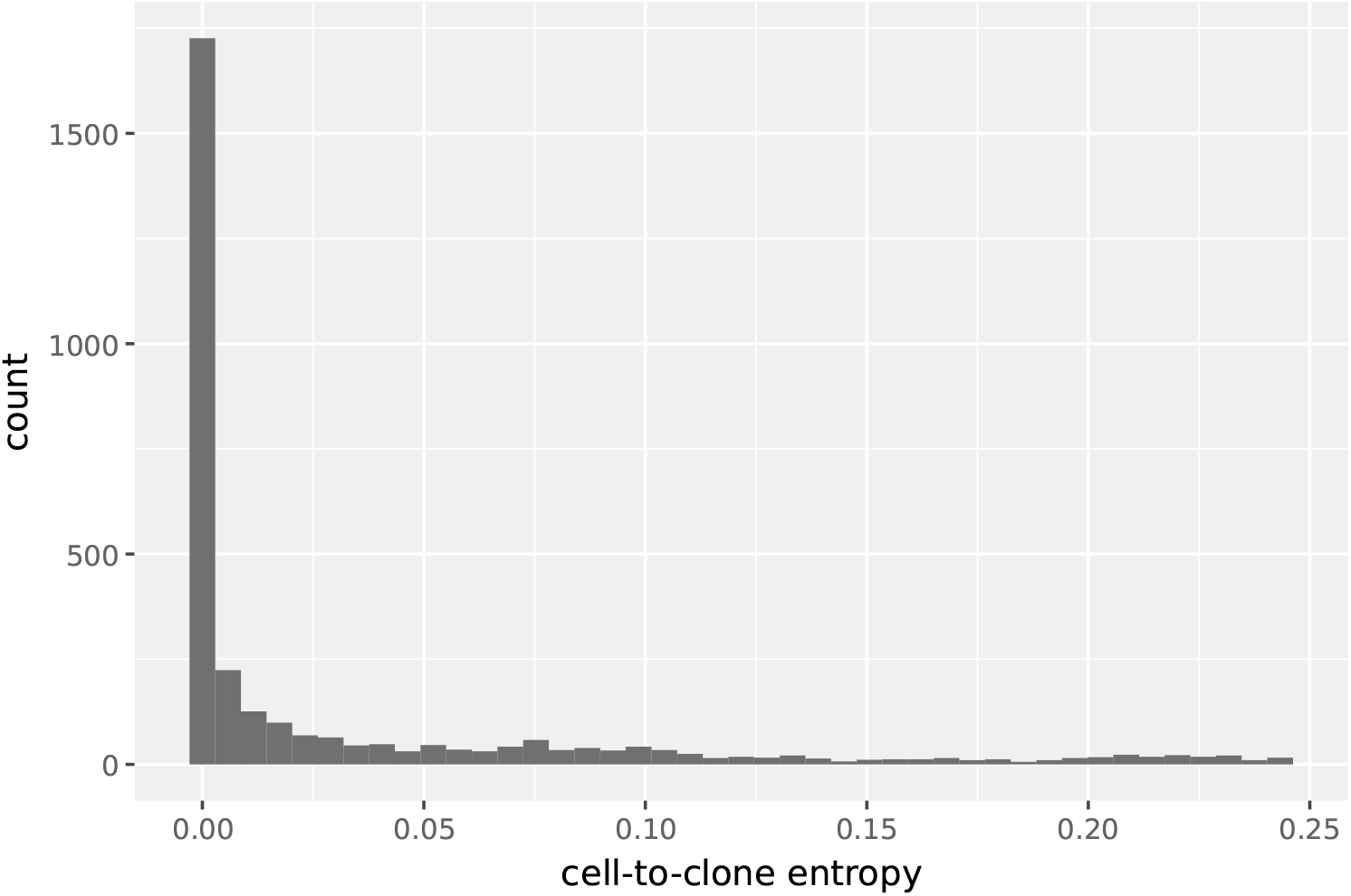
Distribution of cell-to-clone variational entropy corresponding to GEM2.3 clone tree *T*^∗^.

**Figure S9:**
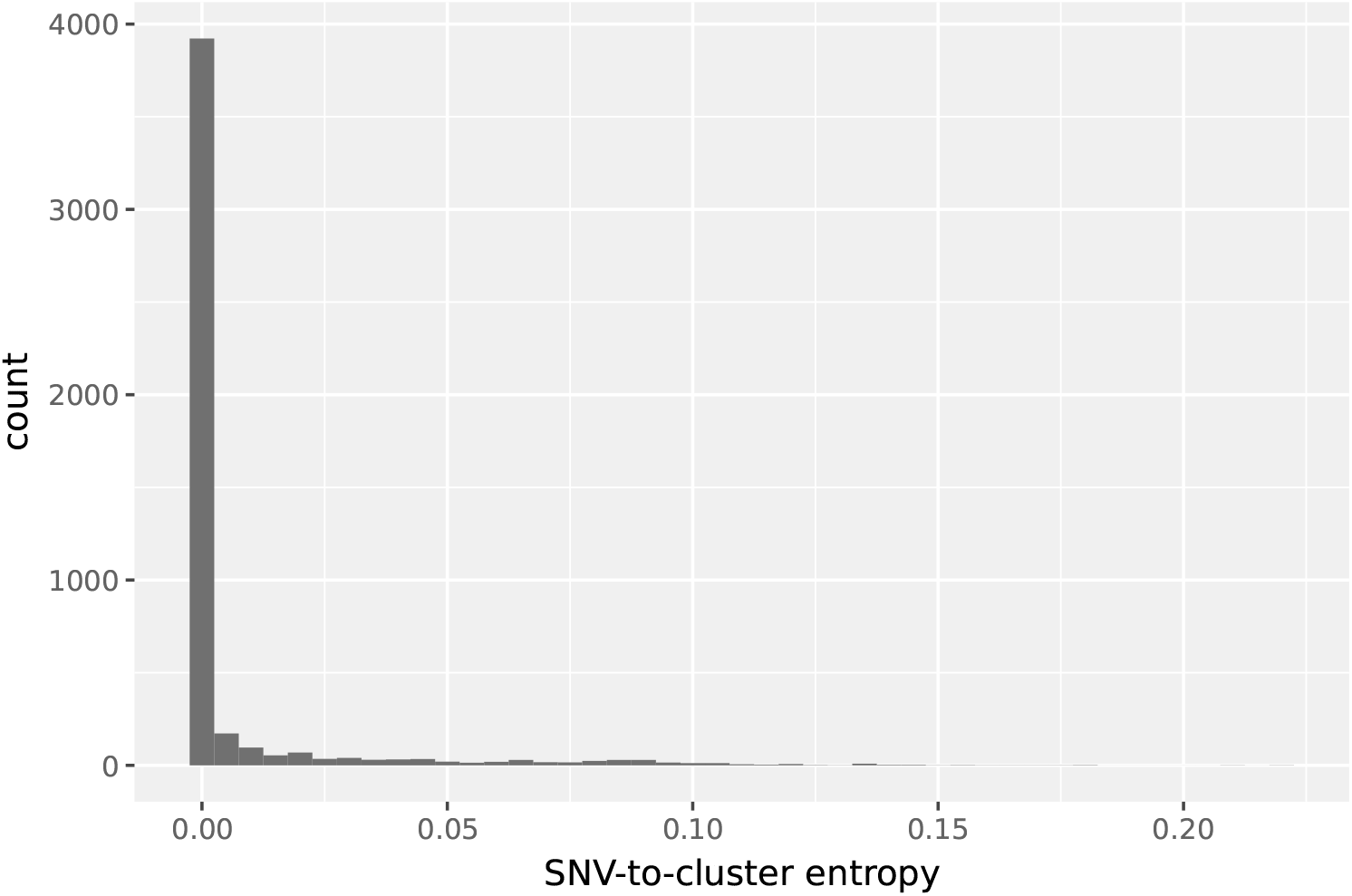
Distribution of SNV-to-cluster variational entropy corresponding to GEM2.3 clone tree *T*^∗^.

**Figure S10:**
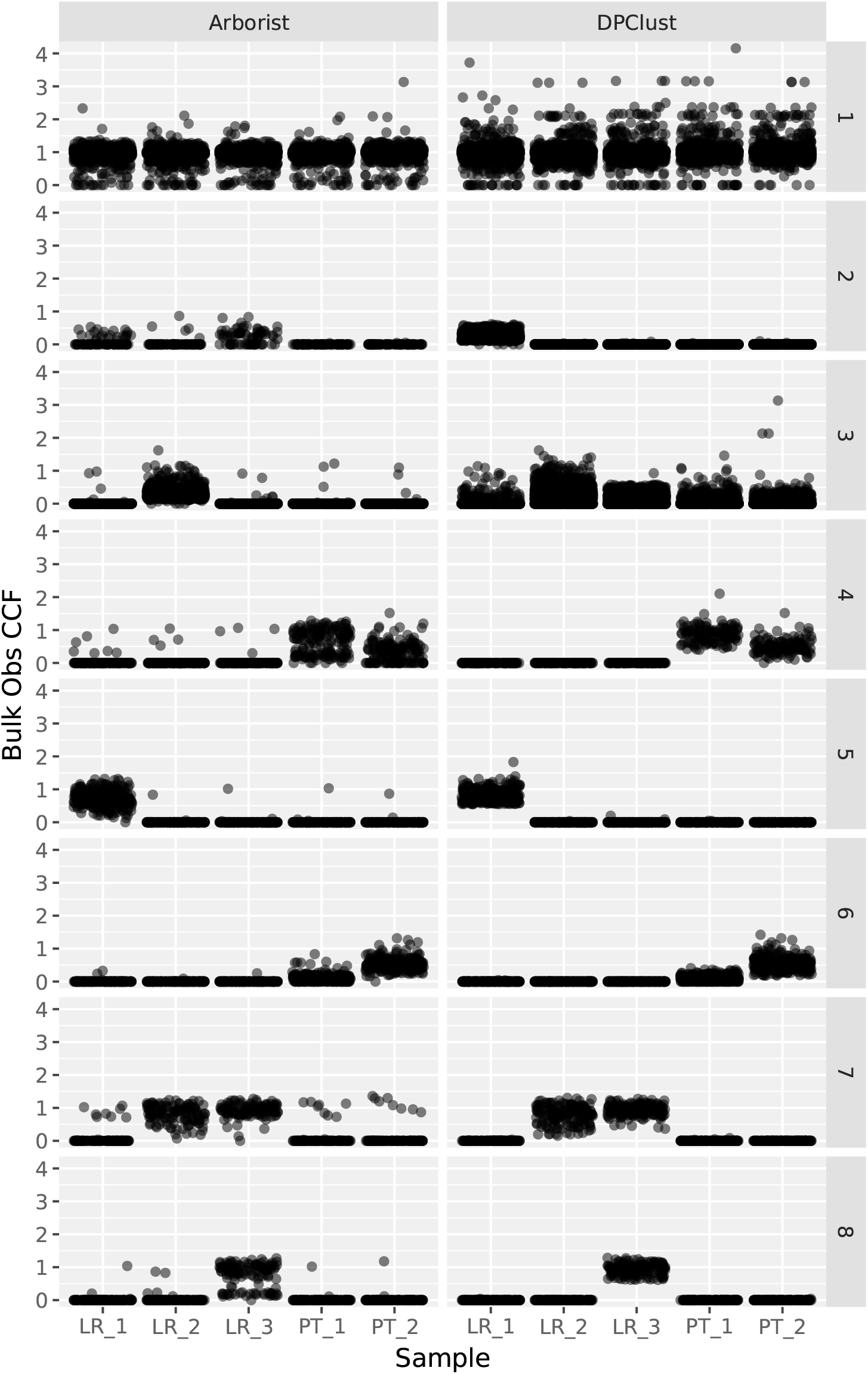
Comparison of bulk sequencing observed CCFs by SNV-to-cluster label and sample corresponding to GEM2.3 clone tree *T*^∗^.

